# Small molecule allosteric inhibitors of RORγt block Th17-dependent inflammation and associated gene expression in vivo

**DOI:** 10.1101/2021.02.19.431952

**Authors:** Steven A. Saenz, Andrea Local, Tiffany Carr, Arvind Shakya, Shivsmriti Koul, Haiqing Hu, Lisa Chourb, Justin Stedman, Jenna Malley, Laura Akullian D’Agostino, Veerabahu Shanmugasundaram, John Malona, Eric Schwartz, Lisa Beebe, Meghan Clements, Ganesh Rajaraman, John Cho, Lan Jiang, Alex Dubrovskiy, Matt Kreilein, Roman Shimanovich, Lawrence G. Hamann, Laure Escoubet, J. Michael Ellis

## Abstract

Retinoic acid receptor-related orphan nuclear receptor (ROR) γt is a member of the RORC nuclear hormone receptor family of transcription factors. RORγt functions as a critical regulator of thymopoiesis and immune responses. RORγt is expressed in multiple immune cell populations including Th17 cells, where its primary function is regulation of immune responses to bacteria and fungi through IL-17A production. However, excessive IL-17A production has been linked to numerous autoimmune diseases. Moreover, Th17 cells have been shown to elicit both pro- and anti-tumor effects. Thus, modulation of the RORγt/IL-17A axis may represent an attractive therapeutic target for the treatment of autoimmune disorders and some cancers. Herein we report the design, synthesis and characterization of three selective allosteric RORγt inhibitors in preclinical models of inflammation and tumor growth. We demonstrate that these compounds can inhibit Th17 differentiation and maintenance *in vitro* and Th17-dependent inflammation and associated gene expression *in vivo*, in a dose-dependent manner. Finally, RORγt inhibitors were shown to inhibit tumor formation in pancreatic ductal adenocarcinoma (PDAC) organoids.

## Introduction

Psoriasis is a chronic, immune-mediated disease characterized by the presence of large, erythematous, scaly plaques commonly found at multiple sites on the skin surface [1-3]. Psoriatic skin lesions display increased infiltrates of multiple lymphocyte lineages, including T helper type 17 (Th17) cells, γδT cells and innate lymphoid cells (ILCs), in the epidermal and dermal layers [3]. In addition, elevated gene expression levels of proinflammatory cytokines including TNFα, IL-17A, IL-22 and IL-23 have been reported in skin biopsies from psoriatic patients [2-4]. These cytokines are known to act on various cell types within the skin tissue microenvironment, including keratinocytes, neutrophils, endothelial cells and fibroblasts which, in turn, promote aberrant keratinocyte activation, hyperproliferation and tissue inflammation. For patients with moderate-to-severe psoriasis, treatment options are limited. Phototherapy or systemic medications including methotrexate and cyclosporine are common, as are neutralizing monoclonal antibodies against TNFα. However, these therapies are not broadly efficacious.

Retinoic acid receptor-related orphan nuclear receptor c (RORC) is a nuclear hormone receptor in the retinoid acid receptor-related orphan receptor (ROR) subfamily of transcription factors including two isoforms that vary at the N-Terminus [5]. RORγ is widely expressed while RORγt is induced during the transition from double negative to double positive thymocytes where it regulates the survival factor Bcl-xL, allowing for maturation into single positive T cells [5, 6]. Beyond its role in thymopoiesis, RORγt is expressed in subsets of immune cells including γδT cells, Th17 cells, ILC3, NKT cells and NK cells [5]. RORγt is the master regulator of Th17 cells. In response to IL-1α/β, IL-6 and IL-23, it regulates differentiation of Th17 cells as well as maintenance and production of cytokines including: IL-17A, IL-17F, IL-22 and granulocyte-macrophage colony stimulating factor (GM-CSF) [7, 8]. The primary function of Th17 cells is to regulate immune responses that lead to clearance of extracellular pathogens including bacteria and fungi. However, excessive IL-17A production has been linked to autoimmune diseases such as psoriasis, psoriatic arthritis, rheumatoid arthritis and multiple sclerosis [9]. Thus, the discovery that RORγt regulates the development of multiple lymphocyte lineages including IL-17A-producing immune cell populations provides compelling evidence that disruption of the RORγt/IL-17A/IL-23 axis may represent a viable therapeutic option for the treatment of psoriasis.

Targeting the RORγt/IL-17A/IL-23 axis, either by genetic manipulation or antibody-mediated neutralization of pathway cytokines (e.g. IL-17A, IL-23 and GM-CSF) ameliorates disease pathology in multiple animal models of autoimmunity and inflammation. These findings extend to patients, where biologic therapies targeting IL-23 and IL-17A or their receptors have demonstrated clinical efficacy in psoriasis, psoriatic arthritis, autoimmune uveitis and ankylosing spondylitis. In addition, small molecules targeting the RORγt/IL-17A/IL-23 axis have demonstrated clinical efficacy through reduction in circulating IL-17A levels. In multiple phase 3 trials in psoriasis patients, Otezla (Apremilast), a phosphodiesterase 4 (PDE4) inhibitor, reduced the production of IL-17A, IL-17F, IL-22 and TNFα by 40-50% at Week 4 with concomitant Psoriasis Area and Severity Index (PASI) -75 response rates of approximately 30% at Week 16 [10]. Furthermore, VTP-43742, an orthosteric RORγt inverse agonist, demonstrated 50-75% reductions in IL-17A and IL-17F levels in the plasma of plaque psoriasis patients with 25-30% PASI-75 response rates at Week 4 [11]. These data demonstrate that small molecule inhibitors targeting RORγt can block Th17-associated protein production across multiple cell populations with improved clinical outcomes.

As a key regulator of CD4^+^ T-cell polarization and Th17 cell function, RORγt is thought to play a key role in tumor immunity [12, 13]. In fact, knockout of RORγt in adult mice leads to development of lymphoblastic lymphomas within 6 months in a manner similar to embryonic RORγt loss [14]. RORγt agonists are currently in clinical trials for multiple indications including NSCLC and ovarian cancer [15]. However, as Th17 cells have been ascribed both pro- and anti-tumor effects, based on disease type and presence of other immune cells and cytokines, the role of RORγt in tumor immunity is controversial [12]. A recent study demonstrated high RORγt expression in more advanced and metastatic pancreatic cancer patients. The study utilized transcriptomic and epigenetic profiling of a pancreatic ductal carcinoma (PDAC) KP/C mouse model to identify transcription factors important for cancer stem cell (CSC) maintenance and growth [16]. PDAC CSCs are known to be resistant to cytotoxic therapies like standard of care gemcitabine, and higher CSC levels are associated with decreased survival [17]. The contribution of several transcription factors to tumor cell growth, including RORγt, was confirmed using CRISPR knockout screens in mouse PDAC organoid models. Furthermore, it was demonstrated that genetic ablation or inverse agonism of RORγt activity, with SR2211, was sufficient to inhibit both human and mouse tumor growth [16].

Given the level of clinical validation for targeting the RORγt/IL-17A/IL-23 pathway, our aim was to evaluate additional allosteric inhibitors, complementary to previously reported ROR antagonists [18, 19], in relevant pre-clinical assays to identify a suitable candidate for further drug development. Herein we detail the design, synthesis and pre-clinical characterization of three selective allosteric RORγt inhibitors, Compounds (Cmpd) 1, 2 and 3, in models of inflammation and tumor growth. Potencies of these compounds were determined using GAL4 reporter assays and human primary cell Th17 differentiation and maintenance assays. Relationships between pharmacokinetics and pharmacodynamics (PK/PD) were established by monitoring Th17-associated gene expression after dosing in Th17-dependent mouse models of imiquimod-induced skin inflammation and experimental autoimmune encephalitis (EAE). Finally, the antitumor activity of Cmpd 3 was assessed in PDAC organoids and the genetically engineered KP/C mouse model.

## Materials and Methods

### Mice & imiquimod (IMQ)-induced skin inflammation & experimental autoimmune encephalomyelitis (EAE) model

For IMQ-induced skin inflammation, wild-type (WT) Balb/c female mice, 5-8 weeks of age, were purchased from Charles River Laboratories. All mice were housed in pathogen-free conditions at Bristol Myers Squibb (Cambridge, MA). Aldara cream containing 5% imiquimod (Patterson Veterinary Supply, Inc; Devens, MA) was applied to the ears daily (11-14 mg) for 3 days. Mice were treated on days 1-4) per os (PO) with vehicle (0.5% methylcellulose/0.25% Tween80) or RORγt inhibitors at indicated doses immediately prior to Aldara application. Ear thickness was measured using micro-calipers. Tissues were harvested on day 4, 2 hours following the last compound dose.

EAE studies were conducted at Hooke Laboratories. WT female C57BL/6 mice (Taconic Labs) 9 weeks of age were inoculated on day 0 with MOG_35-55_ peptide (Hooke Kit™ MOG35-55/CFA Emulsion PTX) (EK-2110, Hooke Laboratories, Lawrence MA) followed by intraperitoneal (IP) injections of pertussis toxin at 2 and 24 hours. Mice were treated twice daily (starting at day 1) per os (PO) with vehicle (0.5% MC/0.25% Tween80) or RORγt inhibitors at indicated doses. FTY720 (Gilenya) was dosed once daily at 3 mg/kg starting at day 1. EAE clinical scores were evaluated daily and scored from 0-5 according to Hooke Lab EAE scoring guidelines http://hookelabs.com/services/cro/eae/MouseEAEscoring.html. Mean clinical scores & body weight loss were assessed and statistical significance calculated by Wilcoxon’s non-parametric or 2-tailed Student’s t-test, respectively. All studies performed were approved in accordance with the Institutional Animal Care and Use Committee of Bristol Myers Squibb and complied according to Bristol Myers Squibb guidelines.

### *In vivo* tumor growth suppression

Pancreatic tumor chunks from KPC mice [20] on C57Bl/6 background were inoculated subcutaneously in the flank of syngeneic mice. Mice were randomized and distributed in 5 groups of 8 mice each with an average tumor volume of 100 mm^3^. Mice were treated with Cmpd 3 (30mg/kg, BID), Gemcitabine (120mg/kg, QW) or Cisplatin (5mg/kg, QW). Tumor volumes and body weight were measured twice each week.

### Pancreatic organoid cell culture

Patient derived organoids were established as previously described [21]. Briefly, PDX tumor chunks were minced then enzymatically digested into single cells by using a tumor dissociation Kit (Miltenyi biotec Cat# 130-095-929). Mouse and human cells were separated through magnetic separation, and isolated tumor cells were cultured on Matrigel-coated dishes. Organoid cultures were maintained by growing on Matrigel in human complete maintenance media: Advanced DMEM/F12 media (Life Technology, Cat# 12634028), 10 mM HEPES (Life Technology, Cat# 15630-080), 1X Glutamax (Life Technology, Cat# 35050-061), 1X Antibiotic-Antimycotic (Life Technology, Cat# 15240-062), 10 mM Nicotinamide (Sigma-Aldrich, Cat# N0636-500G), 250 ng/mL R-Spondin-1 (Peprotech, Cat# 120-38), 100 ng/mL Noggin (Peprotech, Cat# 120-10C), 50 ng/mL EGF (Peprotech, Cat# AF-100-15), 100 ng/mL FGF10 (Peprotech, Cat# 100-26), 10 mM Gastrin-1 (Sigma-Aldrich, Cat# G9020), 500 nM A83-01 (Sigma, Cat# SML0788), 20 μM Y-27632 (LC Laboratories, Cat# Y-5301), 1X B27 (Life Technology, Cat# 12587010), 10 ng/mL Wnt3a (R&D Systems, Cat# 5036-WN-010), 0.1 % Methylcelluose (R&D Systems, Cat# HSC001) and Plasmocin (Invivogen, Cat# Ant-mpp). and were passaged every 10-14 days. Organoids were isolated from Matrigel (Corning, Cat# 08-774-406) in a cell recovery solution (Corning, Cat# CB-40253) for minimum of about two hours. Spheroid clusters were then dissociated into single cell suspension with TrypLE (Gibco, Cat# 12605). After dissociation, single cells were suspended in the complete growth medium as described above. After counting, a single-cell organoid suspension was plated on pre-warmed matrigel coated plates.

### *In vitro* organoid growth assay

Organoids were isolated and dissociated as described above. Cell numbers were counted by trypan blue exclusion and 5000 cells per well were plated on the Matrigel coated and pre-warmed 96-well plates. Compounds (Cmpd 3 and SR2211) were prepared in DMSO to a stock concentration of 10 μM and were added in indicated doses (0.03 μM to 30 μM) either on the first day or third day of plating. 100 μl of CellTiter-Glo® 3D reagent (Promega cat# G9682) was added to each well after desired time points and the luminescence signal was measured after 30 minutes.

### RNA collection, Gene expression, Cytokine production analysis

For RNA isolation & gene expression, ears are collected on day 4 in RNAlater (Qiagen) and stored at 4° C until processed. Ears were homogenized using Procellys 24 homogenizer & hard tissue homogenizing beads (Bertin Instruments), 2 cycles of 30 seconds @ 6000 rpm in RLT lysis buffer according to manufacturer’s instructions and RNA isolated using RNAeasy Plus MiniPrep columns (Qiagen). cDNA is generated using SuperScript VILO cDNA Synthesis kit (Invitrogen) & gene expression assessed using TaqMan Fast Master Mix & and TaqMan FAM-MGB probe sets (Applied Biosystems): *Gapdh*, Mm99999915_g1, *Il17a*, Mm00439618_m1, *Il17f*, Mm00521423_m1, Il22, Mm01226722_g1 & Bclxl, Mm00437783_m1. QPCR reactions were run on QuantStudio 7 instrument. Relative quantification and fold changes were calculated using *dd*CT values against *Gapdh* and normalized to control-treated animals. In tumor systems, RNA was isolated from tumor sections (Qiagen RNeasy kit cat#74106) then cDNA was prepared using 2 μg of RNA and the High capacity RNA to cDNA kit (Applied Biosystems cat#4387406) as per manufacturer’s instructions. Biomarker expression was determined using Taqman gene expression probes (Supplemental Table 1) and Universal master mix (ThermoFisher Scientific cat#4305719) with expression levels normalized to *Gapdh*.

For cytokine production, ears are removed, split in half using forceps and floated, dermis side-down, in DMEM media (Gibco) and incubated at 37° C for 24 hours. Following incubation, media was removed and cytokine production assessed by Luminex assay (Bio-Rad Laboratories).

### Bioanalysis & Pharmacokinetic measurements

In the IMQ-induced inflammation model, whole blood (300-500 μL) was collected and centrifuged (1000 g x 10 min) at 20° C to obtain plasma samples. In KP/C mouse model, tumors were collected and homogenized with phosphate buffer at ratio of 1:3 (w:v). Plasma standard curves were prepared by adding each test compound in to mouse plasma and serial diluting to desired concentration. Blank tumor homogenate and blank plasma was add to plasma standards and tumor homogenate samples, respectively, at 1:1 (v:v) ratio for matrix match of tumor sample analysis. An aliquot of 50 μL of each plasma sample, each tumor sample and each corresponding standards was added to 200 μL of acetonitrile with 100 ng/mL of carbutamide (Sigma Aldrich, St. Louis, MO), internal standard (IS), for protein precipitation, then filtered through a 96-well Orochem filtration plate (Orochem Technologies Inc., Naperville IL). Each extracted test compound in resultant supernatant was analyzed with appropriate liquid chromatography column eluting to a Sciex QTRAP 6500+ LC/MS/MS system (Applied Biosystems, Foster City, CA). Each analyte was characterized by Turbo IonSpray ionization multiple reaction monitoring (MRM). Quantitative drug concentrations were determined by standard calibration curve analysis, using linear fitting with 1/x^2^ weighted plot of the analyte/IS peak area ratio *vs* analyte concentration.

### *In vitro* human Th17 cell differentiation & maintenance cultures

For differentiation assays, naïve CD4^+^ T cells were enriched (StemCell Technologies; 19555) from healthy donor human PBMCs and cultured in 96-well plates, with XVIV0-15 (Lonza; 04418Q) and anti-CD3/anti-CD28 Dynabeads (Thermo; 11161D). Th0 cell cultures were provided IL-2 (15 U/mL; R&D Systems 202-IL-500), while Th17 cell cultures were supplemented with IL-6 (20 ng/mL; R&D Systems 206-IL-010), TGF-β (10 ng/mL; R&D Systems 240-B002), IL-23 (10 ng/mL; R&D Systems 1290-IL-010) and IL-1β (10 ng/mL; R&D Systems; 201-LB-005). Compound or DMSO vehicle control was added on day 0, and on day 6 supernatants and cells were harvested for Luminex and flow cytometry, respectively. Supernatants were analyzed using a human Th17 cell cytokine panel multiplex (BioRad 171AA001M). For flow cytometry, cultured cells were incubated with phorbol 12-myristate 13-aetate (PMA) and ionomycin (eBioscience; 00-4333-57), in the presence of GolgiStop (BD; 554724) for 5 hours at 37° C. Single cell suspensions were stained with a fixable viability dye (Invitrogen; L34966) and intracellular staining for IL-17A (eBioscience; 50-7179-42; eBio64Dec17) and IFNγ (BD; 563563; b27) was performed as described in the Foxp3/Transcription Factor Staining Buffer kit (eBiosience; 00-5523-00). For Th17 cell maintenance cultures, Th17 cells were enriched (StemCell Technologies; 17862) from healthy donor human PBMCs and cultured in 96 well plates in IMDM supplemented with 10% FBS, penicillin (10 U/mL), streptomycin (10 μg/mL), glutamine (2 mM), and β-mercaptoethanol (55 μM) in the presence of anti-CD3/anti-CD28 Dynabeads (Thermo; 11161D). Th17 cell maintenance cultures were supplemented with IL-23 (50 μg/mL) and IL-1β (10 ng/mL). Compound or DMSO vehicle control was added on day 0, and on day 4 supernatants and cells were harvested for Luminex and flow cytometry, respectively, as described above.

### Statistical analyses

Statistical significance was determined using GraphPad Prism 8 Student’s *t*-test or one-way ANOVA with Tukey’s multiple comparisons test as indicated. Data presented are mean ± SEM. A *P*-value equal to or less than 0.05 was considered to be statistically significant.

## Results

### Design, synthesis and characterization of RORγt allosteric antagonists

Three RORγt allosteric antagonists, similar to previously described molecular architectures disclosed by Lycera (footnote: patent estate licensed to Celgene in 2017) and Merck [19], were designed, synthesized, and characterized in a suite of immunology and oncology assays. Design of these antagonists was focused on an indazole core with aims of enhancing ligand efficiency, facilitating synthetic preparation, and improving physicochemical properties. Cmpd 3 emerged as a lead candidate based upon its favorable potency and selectivity, coupled with a moderate oral exposure across species, synthetic accessibility, and physicochemical properties.

A fit for purpose route to access Cmpd 3 was developed, employing 12 total synthetic steps and a longest linear sequence of 7 steps from commercially available starting materials (Figure 1). Cmpd 3 was prepared in 3% yield from 2,6-difluorobenzaldehyde. Acylation of diaholindazole Intermediate 1 (Int-**1**) was followed by a Suzuki –Miyaura coupling with enone Int-**3** in good yield. Treatment of the lithium enolate of Int-**4** with Mander’s reagent gave the keto ester Int-**5**, which was reduced under Noyori conditions to give alcohol Int-**6**. Hydrolysis afforded the acid Int-**7**, which was recrystallized with (R)-(+)-1phenylethylamine to provide stereo-enriched Int-**8** in a 33% yield and 99% purity. Treatment of the Int-**8** saltwith citric acid delivered the final compound 3. Single molecule X-ray crystallographic analysis confirmed the stereochemical configuration of Cmpd 3 as (*R,R*) (Figure 2). Access to Cmpd 1 and Cmpd 2 was accomplished in similar fashion.

**Figure 1:**
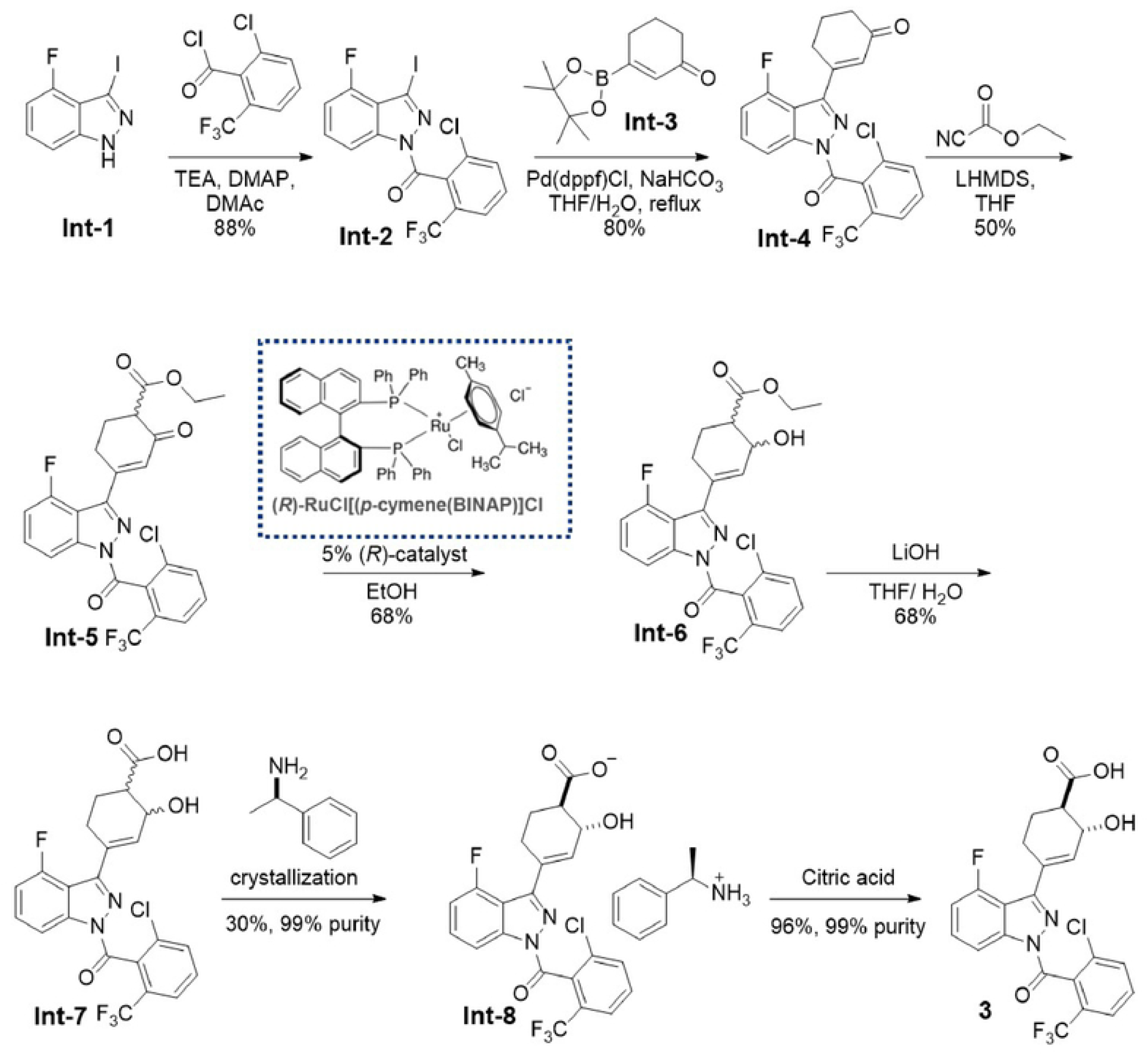
Representative synthesis of Cmpd 3, an allosteric RORγt inhibitor

**Figure 2:**
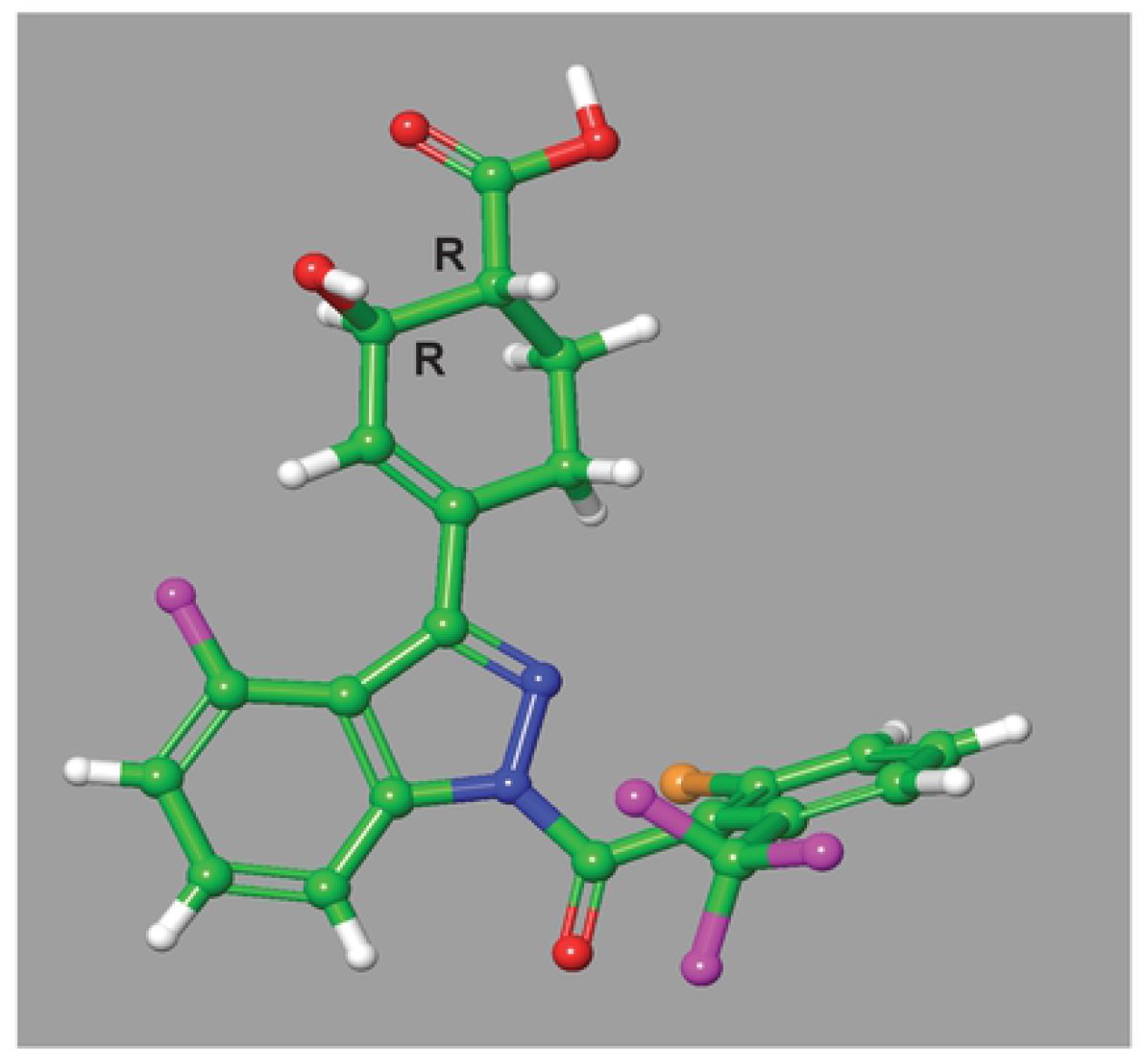
Single molecule X-ray crystallography of Cmpd 3. Small molecule X-ray structure of Cmpd 3 indicates only the (*R,R*)-enantiomer is present in the crystal structure. Crystal conformation of compound is and depicted in ball and stick representation (carbon – green; hydrogen – white; oxygen – red; nitrogen – blue; fluorine – magenta; chlorine – orange).

#### RORγt inhibitors attenuate human Th17 cell differentiation and maintenance

Using a GAL4 luciferase reporter assay consisting of the ligand binding domain of either human or mouse RORγt fused to the DNA binding domain of GAL4, nanomolar (nM) cell-based IC_50_’s were observed for Cmpd 1, Cmpd 2 and Cmpd 3 (data not shown). To further profile these inhibitors in a more biologically-relevant assay, the impact of treatment with RORγt inhibitors Cmpd 1, Cmpd 2 and Cmpd 3 on Th17 cell differentiation was assessed *in vitro*. RORγt is the central transcriptional regulator of Th17 cell identity, promoting expression of key subset effectors, including the lineage defining cytokine IL-17A [22, 23]. Previous studies have shown, through genetic deletion or small molecule inhibition, that RORγt is crucial for the development of Th17 cells and contributes to the maintenance of Th17 cell function [22, 24-29]. Human naïve CD4^+^ T cells were cultured under Th17 cell polarizing conditions in the presence of titrating doses of compounds (Figure 3A). All three RORγt inhibitors blocked IL-17A secretion in a dose dependent manner with approximately 95% maximal inhibition relative to DMSO vehicle control (Figure 3D). All compounds had single digit nanomolar IC_50_’s (Supplementary Table 2), with no overt cytotoxicity (Supplementary Figure 1A and B). Intracellular cytokine staining also showed near complete inhibition of Th17 cell polarization, with the percentage of IL-17A^+^ cells returning to levels comparable to those measured in nonpolarizing Th0 cell conditions (Figure 3C). All three RORγt inhibitors were also profiled in human memory Th17 cell cultures, in the presence of lineage maintenance cytokines IL-23 and IL-1β (Figure 3B and E). Again, all compounds reduced IL-17A secretion in a dose dependent manner, with similar nanomolar IC_50_ values and no overt cytotoxicity. Residual IL-17A production by memory Th17 cells is presumably attributable to amplification of the cytokine by additional transcription factors known to regulate its expression. Taken together, Cmpd 1, Cmpd 2 and Cmpd 3 resulted in inhibition of human Th17 cell differentiation and memory Th17 cell IL-17A production *in vitro*.

**Figure 3.**
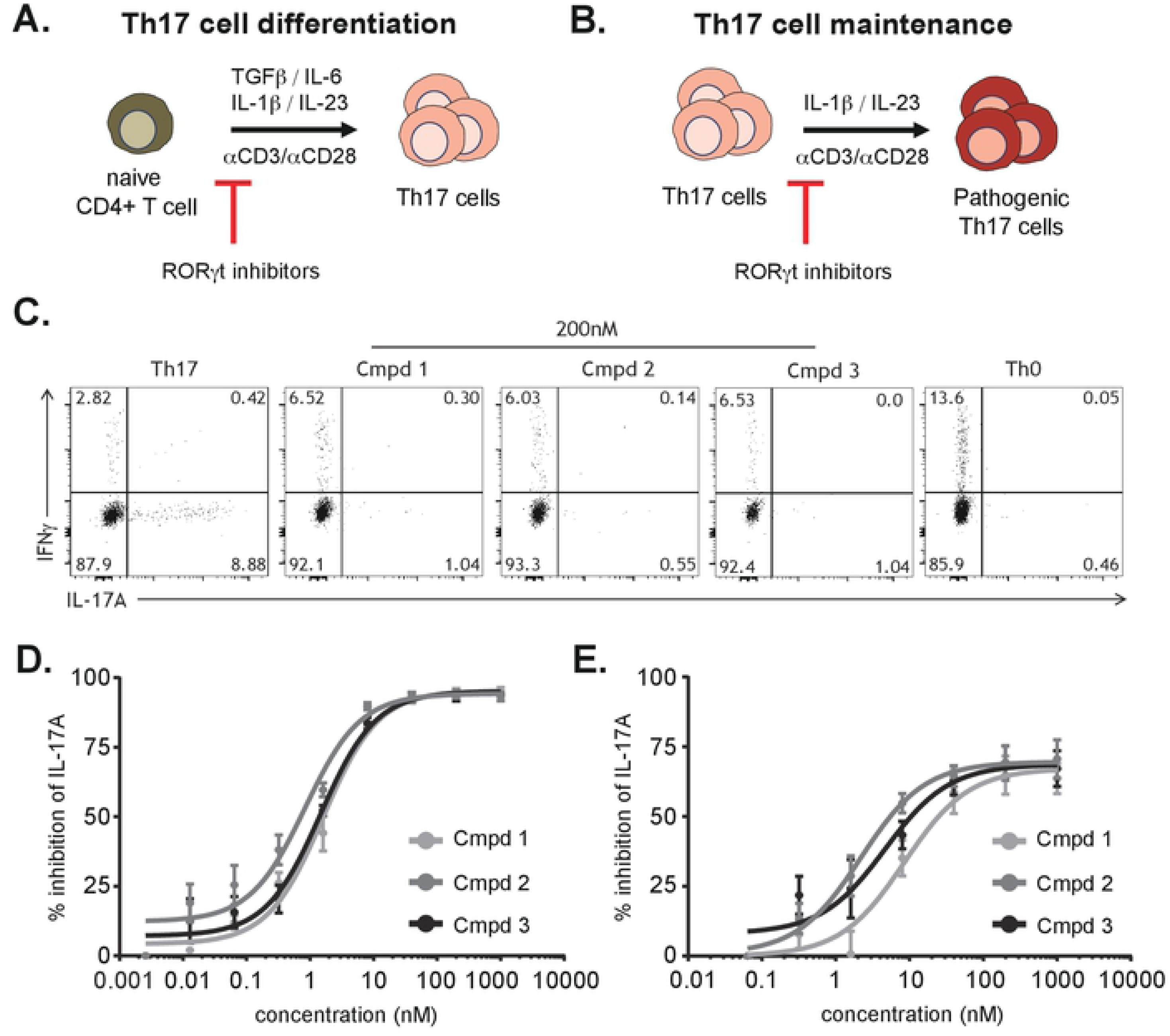
RORγt-mediated inhibition of IL-17A production in human Th17 cell cultures. Schematic of human Th17 differentiation **(A)** or Th17 maintenance **(B)** assays. Representative flow cytometry plots of intracellular cytokine staining from cultures from (A) including Th0 cell nonpolarizing condition **(C)**. IL-17A cytokine production from enriched naïve CD4^+^ T cells from healthy donor PBMC cultured under Th17 cell conditions for 6 days **(D)** or from enriched human Th17 cells from healthy donor PBMC cultured with IL-23 and IL-1β, for 4 days **(E)**, in the presence of the indicated RORγt inhibitors Cmpd 1, Cmpd 2 or Cmpd 3. Data are normalized to and represented as percent inhibition compared to DMSO controls. Error bars are representative of 4 individual donors, from 2 independent experiments.

**Supplementary Figure 1. RORγt-mediated inhibition of IL-17A production in human Th17 cell cultures**. Cell viability of enriched naïve CD4^+^ T cells from healthy donor PBMC cultured under Th17 cell conditions, for 6 days (differentiation) **(A)** or enriched human Th17 cells from healthy donor PBMC cultured with IL-23 and IL-1β, for 4 days (maintenance) **(B)** in the presence of RORγt inhibitors Cmpd 1, Cmpd 2 or Cmpd 3. Data are normalized to and represented as percent of DMSO control. Error bars are representative of 4 individual donors, from 2 independent experiments.

**Supplementary Table 2. RORγt antagonists IC**_**50**_ **values for inhibition of IL-17A production in human Th17 cell cultures**. Concentrations of RORγt inhibitors Cmpd 1, Cmpd 2 or Cmpd 3 in which 50% (IC_50_) IL-17A cytokine was inhibited in Th17 differentiation or Th17 maintenance assays.

#### RORγt inhibitors attenuate imiquimod-induced skin inflammation & Th17-cytokine gene expression

To determine if IC_50_ vs IC_90_ coverage *in vivo* is required to significantly reduce IL-17-dependent gene expression, a 4-day model of Th17-dependent skin inflammation was developed by modifying the pre-clinical model of imiquimod (IMQ)-induced psoriasis. Topical administration of Aldara cream (5% IMQ) is a well-characterized model of Th17 cytokine-dependent skin inflammation [30-32] and a system in which RORγt inhibitors have been shown to attenuate inflammation [29]. Aldara cream was applied daily to the ears of Balb/c mice for 3 days, which were treated daily PO with vehicle or 30, 45 or 75 mg/kg of Cmpd 1. On day 4, ear thickness was measured via calipers and then ears collected for either assessment of Th17-cytokine gene expression or cytokine production. As expected, IMQ-treated animals showed a significant thickening of the ear (0.21 ± 0.03 mm) compared to the control-treated group (0.15 ± 0.006 mm) (Figure 4A). In addition, RNA analysis from skin tissue of IMQ-treated animals demonstrated significantly increased expression of Th17-associated genes, *Il17a, Il17f* and *Il22* as well as IL-17A cytokine production (Figure 4B-D and Supplemental Figure 2), compared to control treated animals. Compared to vehicle treated animals, Cmpd 1 significantly reduced ear thickening at doses of 45 and 75 mg/kg (0.17 ± 0.02 mm and 0.16 ± 0.008 mm, respectively) (Figure 4A), and reduced Th17-cytokine gene expression at all dose levels tested (Figure 4B-D). To determine PK/PD relationships, unbound murine IC_50_ and IC_90_ values were calculated based on human Th17 differentiation IC_50_ & IC_90_ values and adjusted with murine plasma protein binding (data not shown), this extrapolated total plasma concentrations of ∼100 nM and ∼1 μM, which provided free drug level to cover IC_50_ and IC_90_, respectively. Plasma concentration of Cmpd 1 was monitored at 0.5, 2, 4 and 8 hours post-dosing on day 4. The duration of free IC_50_ coverage in plasma, post PO doses of 30, 45 and 75 mg/kg, was 1.8, 2.9 and 3.9 hours, respectively (Figure 4E). In addition to attenuation of Th17-associated gene expression, treatment with Cmpd 1 also reduced IMQ-induced IL-17A cytokine production in ear tissues (Supplemental Figure 2).

**Figure 4.**
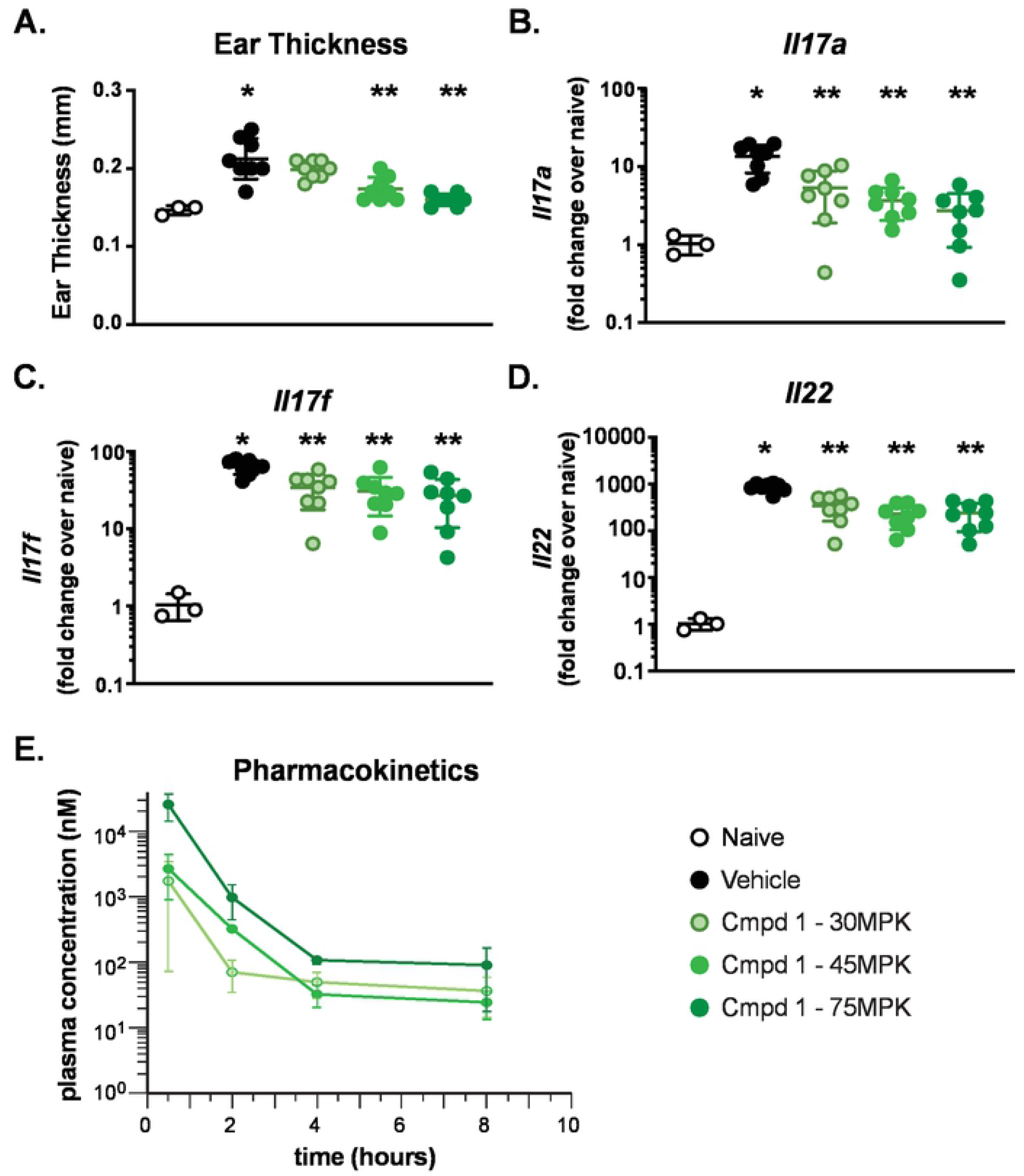
Imiquimod-induced skin inflammation is attenuated by RORγt antagonist Cmpd 1. Ear thickness (mm) was measured in naïve or IMQ-treated animals on day 4 using digital micro-calipers **(A)**. Th17 cytokine gene expression analysis was performed for *Il17a* **(B)**, *Il17f* **(C)** and *Il22* **(D)** on day 4. Expression is normalized to *Gapdh* and presented as fold change over naïve. Kinetic assessment of plasma concentration of Cmpd 1 **(E)**. Each symbol represents an individual animal and error bars denote mean ± SEM. Statistical significance (\**p* ≤ 0.05) was determined using one-way ANOVA with Tukey’s multiple comparisons test, **\***significant over naïve; **\*\***significance over vehicle-treated group. Data are representative of 2 independent experiments with n=3-8/group.

Similar to Cmpd 1, oral administration of indazole-containing RORγt inhibitors, Cmpd 2 and Cmpd 3, resulted in decreased IMQ-induced skin inflammation. Administration of Cmpd 3 at 25, 50 or 100 mg/kg corresponded with unbound IC_50_ coverage of ∼10, 12 and 18 hours and significantly reduced IMQ-induced ear thickening was observed at all doses (0.163 ± 0.002 mm, 0.156 ± 0.001 mm & 0.143 ± 0.002 mm at 25, 50 and 100 mg/kg respectively, compared to 0.17 ± 0.002 mm in control-treated mice). Th17-cytokine gene expression was reduced in the 50 and 100 mg/kg dosed groups (Figure 5A-C). Attenuation of Th17 cytokine responses was observed with unbound IC_50_ coverage of ∼18 hours in plasma, respectively (Figure 5E). Similarly, oral administration of Cmpd 2 resulted in inhibition of Th17-dependent gene expression and inflammation (Supplementary Figure 3). RORγt can also impact *Bclxl* expression in T cell populations, specifically in the thymus [33, 34]. To assess whether *Bclxl* expression was altered by RORγt inhibition in the IMQ-induced skin inflammation, *Bclxl* expression was measured after oral administration of Cmpd 3. Similar to *Il17a* and *Il17f* expression, treatment with Cmpd 3 significantly reduced *Bclxl* expression in skin tissues (Figure 4D).

**Figure 5.**
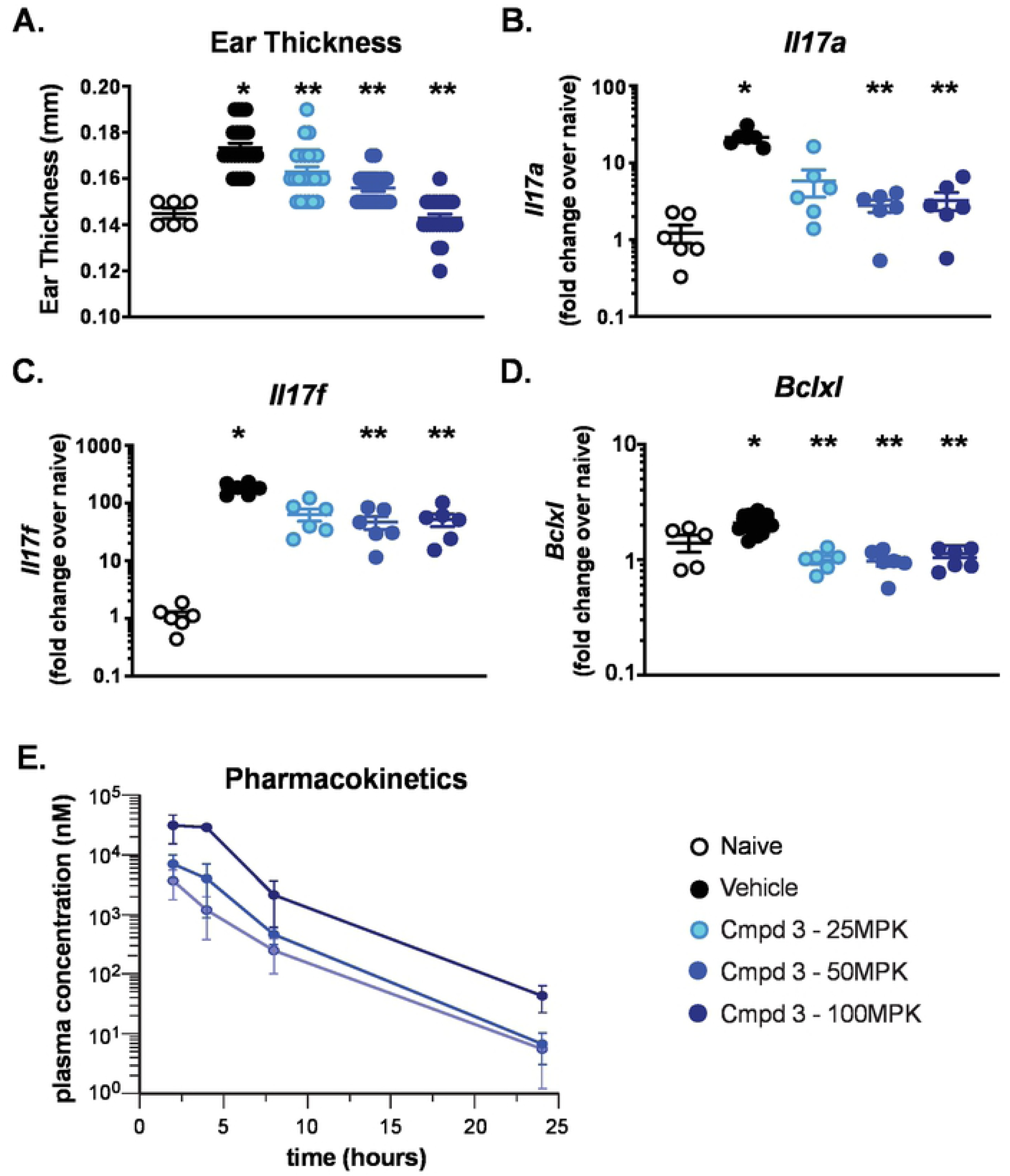
IMQ-induced skin inflammation and RORc activity is inhibited by Cmpd 3. Ear thickness (mm) was measured in naïve or IMQ-treated animals on day 4 using digital micro-calipers **(A)**. Th17 cytokine gene expression analysis was performed for *Il17a* **(B)**, *Il17f* **(C)** and *Bclxl* **(D)** on day 4. Expression is normalized to *Gapdh* and presented as fold change over naïve. Kinetic assessment of plasma concentration of Cmpd 3 **(E)**. Each symbol represents an individual animal and error bars denote mean ± SEM. Statistical significance (\**p* ≤ 0.05) was determined using one-way ANOVA with Tukey’s multiple comparisons test, **\***significant over naïve; **\*\***significance over vehicle-treated group. Data are representative of 1-4 independent experiments with n=6-24/group.

To further assess the coverage required for inhibition of Th17-dependent skin inflammation, ratios of free drug concentrations in plasma and *Il17a* or *Il17f* expressions levels for individual animals were compared. As shown in Figure 6, the concentration of Cmpd 3 in plasma concentration inversely correlated with inhibition of Th17-associated *Il17a* (Figure 6A) and *Il17f* (Figure 6B) gene expression with *r*^*2*^ = 0.37 and 0.44, respectively.

**Figure 6.**
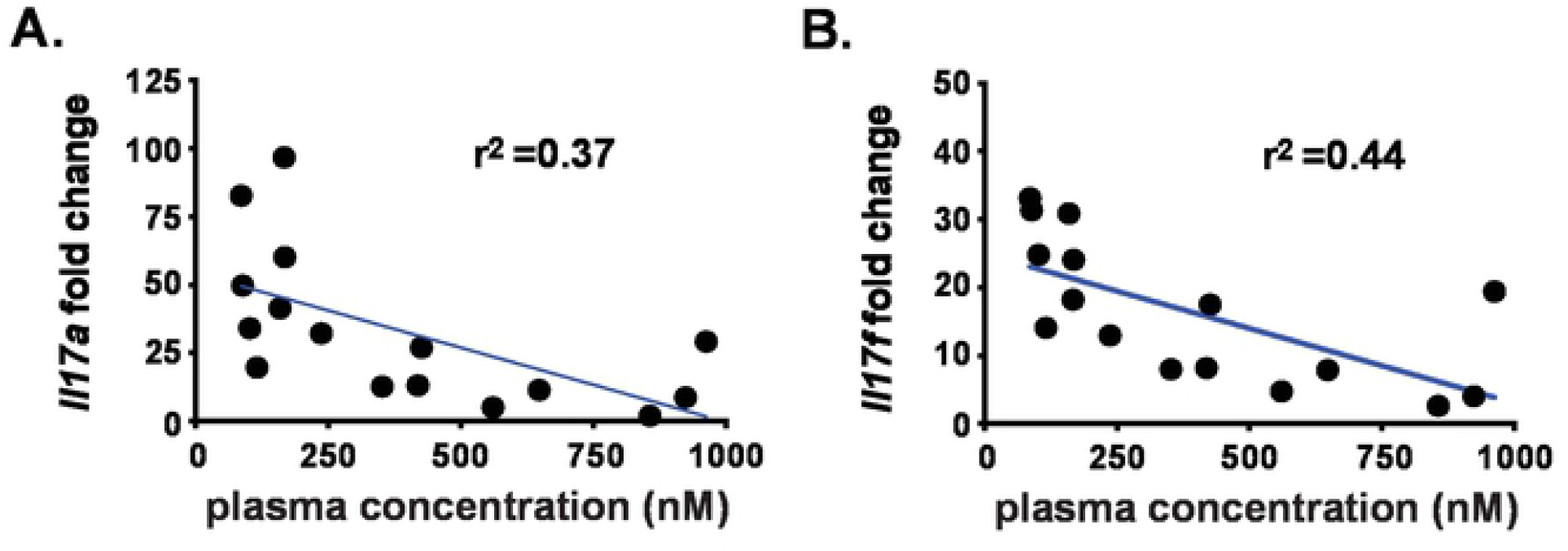
Cmpd 3 plasma concentration inversely correlates with Th17 cytokine gene expression. Th17 cytokine gene expression analysis was performed for *Il17a* **(A)** and *Il17f* **(B)** from IMQ-treated animals dosed with 6 or 60 mg/kg Cmpd 3. Gene expression is normalized to *Gapdh* and presented as fold change over naïve and plotted relative to plasma concentrations for Cmpd 3. Each symbol represents an individual animal and a linear regression (blue line) coefficient was calculated. Statistical significance (\**p* ≤ 0.05) was determined using linear regression analysis. Data are from a single experiment with n=8/group.

**Supplementary Figure 2. RORγt antagonist inhibits IL-17A production in IMQ-treated ear tissue**.

IL-17A cytokine levels were measured by Luminex assay from the supernatants of ear tissue ‘floats’ cultured for 24 hours *ex vivo*. Each symbol represents an individual animal and error bars denote mean ± SEM. Statistical significance (\**p* ≤ 0.05) was determined using one-way ANOVA with Tukey’s multiple comparisons test, **\***significant over naïve; **\*\***significance over vehicle-treated group. Data are representative of 2 independent experiments with n=3-8/group.

**Supplementary Figure 3. RORγt antagonist Cmpd 2 reduces IMQ-induced Th17 cytokine-dependent inflammation**

Ear thickness (mm) was measured in naïve or IMQ-treated animals on day 4 using digital micro-calipers **(A)**. Th17 cytokine gene expression analysis was performed for *Il17a* **(B)**, *Il17f* **(C)** and *Bclxl* **(D)** on day 4. Expression is normalized to *Gapdh* and presented as fold change over naïve. Each symbol represents an individual animal and error bars denote mean ± SEM. Statistical significance (\**p* ≤ 0.05) was determined using one-way ANOVA with Tukey’s multiple comparisons test, **\***significant over naïve; **\*\***significance over vehicle-treated group. Data are representative of 2 independent experiments with n=8/group.

#### RORγt inhibitors reduce the severity of Th17-dependent inflammation in the central nervous system

We next sought to extend these findings to a chronic disease model where we could assess efficacy with prolonged compound exposure. To this end, an EAE model of IL-17-dependent CNS inflammation was employed. Following EAE induction, vehicle-treated animals showed significant disease starting at day 12 and reached a peak clinical score around day 26 of 3.08 ± 0.28 (Figure 7). As a positive control, FTY720 (3 mg/kg) significantly impacted disease onset (day 20) and severity (clinical score 0.54 ± 0.234). Treatment with Cmpd 3 at 3 or 10 mg/kg did not reduce EAE severity, with clinical scores of 2.79 ± 0.25 and 2.32 ± 0.21, respectively. However, Cmpd 3 dosed at 30 mg/kg significantly attenuated EAE severity (clinical score 1.5 ± 0.27) and delayed significant disease onset until day 15 (Figure 7). In addition, inhibition of RORγt led to reductions in Th17-associated gene expression (*Il17a* and *Il22*) in spinal cords and decreased body weight loss in treated-animals compared to vehicle-treated animals (data not shown). These data demonstrate that inhibition of RORγt can provide efficacy in a disease relevant chronic inflammatory model and are consistent with the observation that prolonged duration of IC_50_ coverage *in vivo* results in a more pronounced anti-inflammatory response.

**Figure 7.**
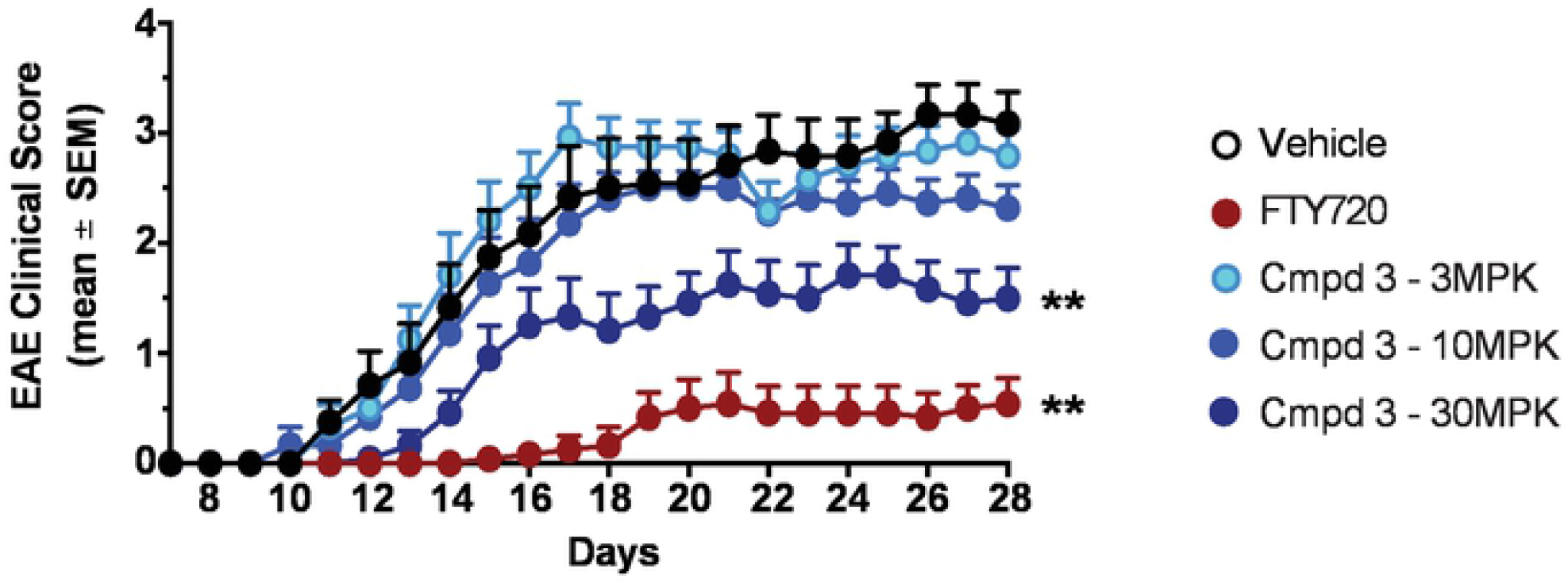
RORγt antagonist protects in EAE immunization model. C57BL/6 mice were immunized with MOG_35-55_ peptide on day 1 and were treated orally, twice daily (from day 1-day 28) with vehicle (0.5% MC/0.25% Tween80) or RORγt inhibitors at indicated doses. FTY720 (Gilenya) was dosed once daily at 3 mg/kg starting at day 1 in MilliQ water. The clinical score (0-5) was determined in these mice from day 7-28. Scores were calculated as follows: 0-no clinical signs, 0.5-partial tail weakness, 1.0-complete tail paralysis, 1.5-flaccid tail & abnormal gait, 2.0-flaccid tail and clear weakness of hind legs, 2.5-partial paralysis in one hind limb, 3.0-complete paralysis in both hindlimbs, 4.0-partial weakness in forelimbs, 5.0-complete paralysis in both forelimbs and hindlimbs). Data are presented as mean ± SEM. Statistical significance of clinical scores & body weight loss were calculated by Wilcoxon’s non-parametric or 2-tailed Student’s t-test, respectively; \**p* < 0.05, compared with vehicle-treated EAE mice. Data are from a single experiment with n=12/group.

#### RORγt antagonism and pancreatic organoid growth

Given the recent finding that identified RORγt as a major regulator in human pancreatic stem cell growth and that the pharmacologic inhibition of RORγt reduced tumor burden [16], we tested the impact of Cmpd 3 as a therapeutic agent for treatment of pancreatic cancer. Patient-derived organoids (PDOs), which have been shown to effectively parallel patient responses to new therapeutic agents [35], were utilized to assay Cmpd 3 activity *in vitro*. In two distinct PDO models, T020P and T031P, compound was added to cultures at either at day 1 or day 3 after plating and organoid formation or growth were evaluated (Supplementary Figure 4). Organoid growth was assayed by CTG at the time points indicated and compared with vehicle treated samples. The RORγt inverse agonist SR2211 [36] inhibited the growth in both PDO models in a time and dose dependent manner with an IC_50_ ∼3 μM. Contrary to this Cmpd 3 decreased the growth of T031P ∼30% at 30 μM after 120 hours of treatment but did not decrease viability of T020P PDOs. Interestingly, Abraxane resistant T020P was also less responsive to treatment with SR2211 (Figure 8A). We compared a single 120 hr treatment with subsequent dosing at 72 hours to rule out drug instability as the reason for lack of efficacy of Cmpd 3 (Supplementary Figure 4). Re-treatment resulted in a slight increase in SR2211 activity but did not boost Cmpd 3 efficacy (Supplementary Figure 5A). Pharmacological blockage of RORγt by Cmpd 3 elicited a modest effect on organoid formation which increased in a time dependent manner for T031P with an IC_50_ of 30 μM at 120 hours and 10 μM at 168 hours. Whereas SR2211 demonstrated greater potency than Cmpd 3 with IC_50_ values ∼1 μM (Figure 8B). In contrast to Cmpd 3, a time dependent increase in activity for SR2211 was observed for all treatment intervals tested (Figure 8 and Supplementary Figure 5). In summary, inverse agonism with SR221 was more effective at decreasing both organoid formation and growth in these systems suggesting that mode of inhibition can effect outcomes in PDAC organoids. Interestingly, the Abraxane resistant PDO model T020P was less sensitive to both compounds (Figure 8 and Supplementary Figure 5), suggesting a common resistance mechanism that may render RORγt inhibition less effective in patients pre-treated with Abraxane. PDO models do not fully recapitulate the heterogeneity and complexity of pancreatic cancer and tumor microenvironment signaling which could explain the lack of activity observed. Therefore, we sought to evaluate the antitumor activity of Cmpd 3 in a genetically engineered mouse model of pancreatic cancer.

**Figure 8.**
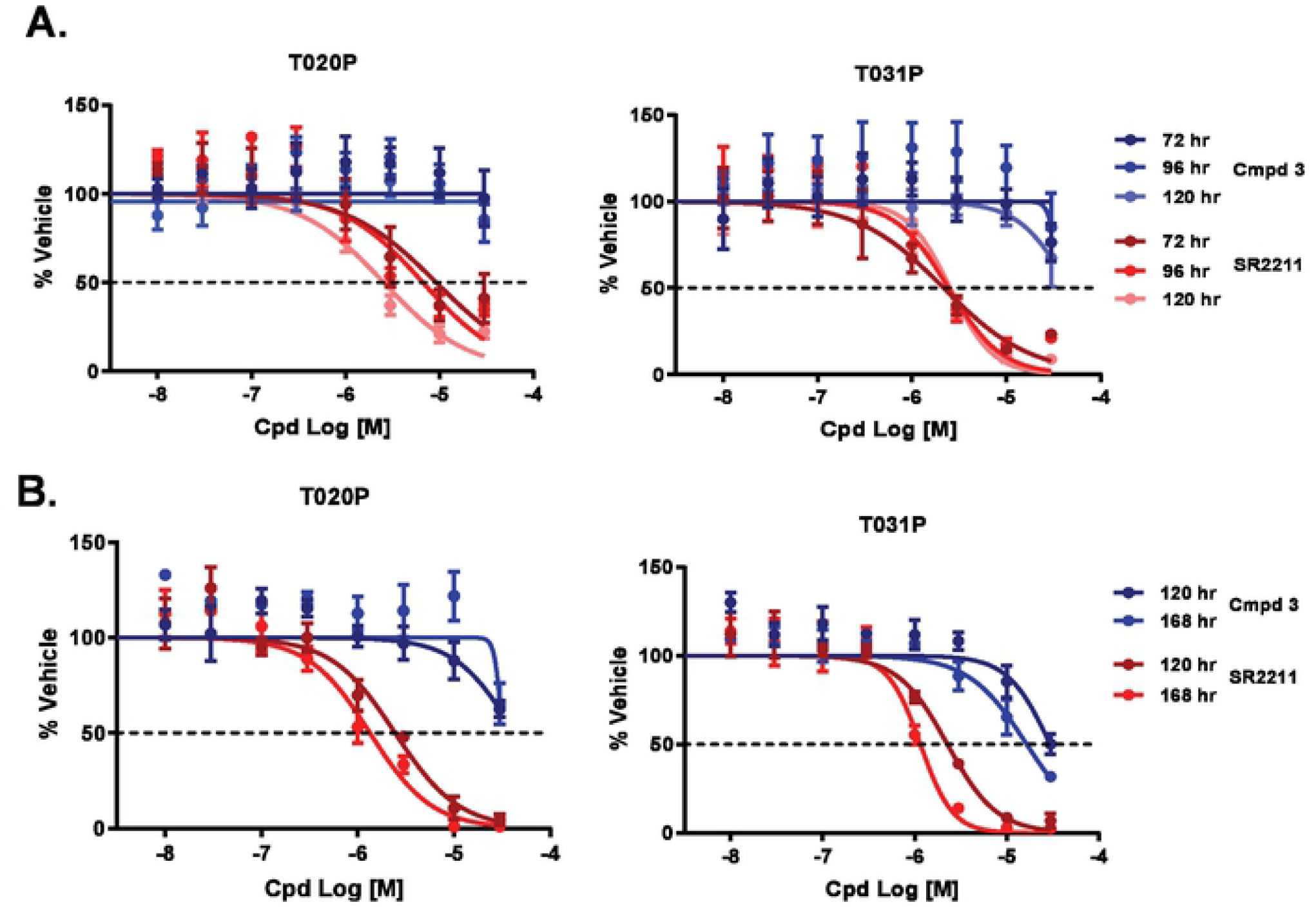
*In vitro* RORγt antagonist activity on PDO growth & formation. PDOs T020P and T031P organoids were dissociated and treated with antagonist (Cmpd 3) or inverse agonist (SR2211) and either organoid growth or organoid formation defects examined. **(A)** Organoid growth assays, PDOs were treated with Cmpd 3 (red line) or SR2211 (blue line) beginning on day 3 after plating for the duration and dose indicated. All values were calculated as % relative to vehicle treated organoids in 3 replicate experiments. **(B)** Organoid formation assays, PDOs were treated with Cmpd 3 (red line) or SR2211 (blue line) starting on day 1 after plating for the duration and dose indicated. All values were calculated as % of vehicle treated organoids in 3 replicate experiments.

**Supplementary Figure 4. Schematic of Patient-derived organoids (PDO) growth & formation assays**. PDOs were dissociated into single cells and plated on day 0. Organoids were treated beginning on day 1 or day 3 to assess effect on organoid formation or organoid growth, respectively. After the indicated treatment schedule organoid growth or formation was assessed by CTG assay.

**Supplementary Figure 5. *In vitro* RORγt antagonist activity on PDO growth & formation. (A)** PDOs were treated with Cmpd 3 or SR221 for 120 hours starting on day 1 after plating. PDOs were either subjected to a single treatment (NR) or compounds were replenished after 72 hours (RT). All values were calculated as % of vehicle treated organoids in 3 replicate experiments. **(B)** PDOs were treated with either Cmpd 3 or SR2211 for 72 or 120 hours starting on day 1 after plating. All values calculated as % vehicle treated organoids in 3 replicate experiments. **(C)** PDOs were treated for 120 hours with Cmpd 3 or SR2211 beginning on day 1 (D1) or day 3 (D3) post-plating. All values were calculated as % of vehicle treated organoids in 3 replicate experiments.

#### Efficacy of Cmpd 3 in KP/C mouse model

To test the hypothesis that inhibition of RORγt can lead to tumor growth inhibition we utilized the Kras^G12D/+^/Trp53^null^/Pdx1-cre (KPC) mouse model of pancreatic cancer model [20]. C57Bl/6 mice were inoculated in the flank with KPC tumor chunks and enrolled in the study when the average tumor volume reached 200 mm^3^. Effect of treatment with Cmpd 3 on tumor growth was evaluated either alone or in combination with gemcitabine and compared to either vehicle or cisplatin treated mice. As expected, gemcitabine and cisplatin significantly reduced the tumor volumes (Figure 9A). However, while Cmpd 3 caused a small decrease in tumor volume the difference was not statistically significant. Additionally, Cmpd 3 conferred no additional benefit when used in combination with gemcitabine (Figure 9A). At the conclusion of the study, tumors were harvested 16 hours post dosing and processed to determine intra-tumoral concentrations of Cmpd 3. Tumor concentrations of Cmpd 3 were highly variable at 47 pmol/g ± 30 and 222 pmol/g ± 251 for Cmpd 3 as a single agent or in combination with gemcitabine, respectively (Figure 8B). Previous PK/PD analysis indicated that these concentrations were sufficient to achieve RORγt antagonism in plasma and skin. To evaluate target engagement in tumors, we examined changes in expression of a subset of potential RORγt target genes in tumor samples. *Msi2* is proposed to be a marker of pancreatic cancer stem cells and a target gene of RORγt [16]. Interestingly, we observed a significant decrease in *Msi2* expression in tumors treated with a combination of Cmpd 3 and gemcitabine but not Cmpd 3 alone (Figure 9C). A similar pattern was observed for putative RORγt target genes *Ehf* and *Ncor2*. However, expression of other putative target genes, *Klf7* and *Osmr*, was decreased in all treatment groups (Figure 9C and Supplementary Figure 6). To summarize, modulation of a variety of RORγt target genes was achieved upon treatment with RORγt antagonist Cmpd 3, alone or in combination with standard of care agent gemcitabine. However, this activity was not sufficient to delay tumor volume in a KPC human tumor mouse model of pancreatic cancer.

**Figure 9:**
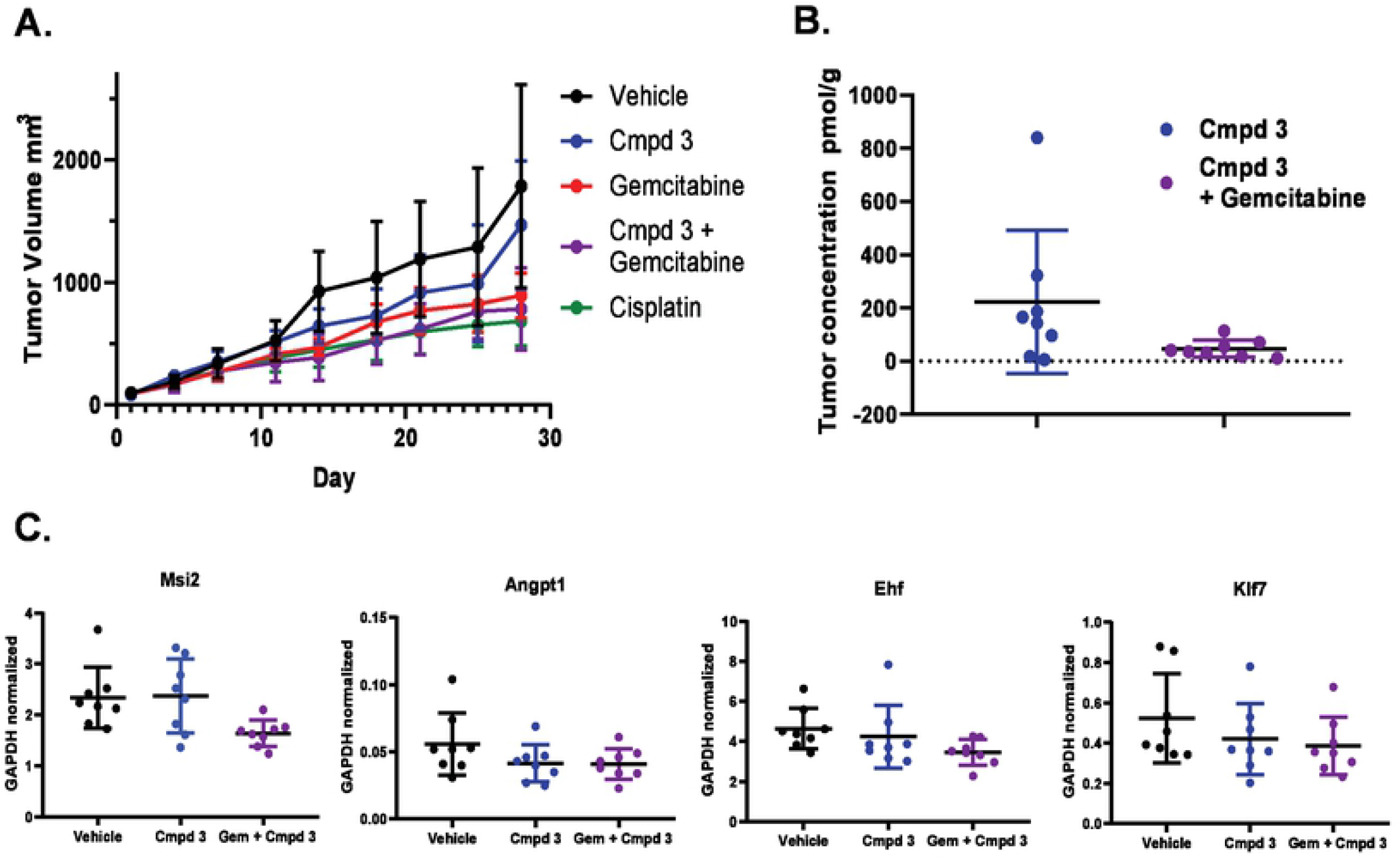
*In vivo* analysis of RORγt antagonist activity on tumor growth. Tumors were implanted in the flanks of C57Bl/6 mice and treatment began when average tumor volume reached 200 mm^3^. **(A)** Tumor volumes from animals treated with vehicle, Cmpd 3 alone, gemcitabine, Cmpd 3 plus gemcitabine or a positive control, cisplatin, were evaluated over 28 days of dosing. Averages were calculated from n=8 mice per group. Error bars represent SD. **(B)** Concentrations were measured by HPLC and plotted as mean ± SD. **(C)** qPCR expression analysis normalized to β-actin expression for vehicle, Cmpd 3, or Cmpd 3 plus gemcitabine treated tumor samples and plotted as mean ± SD.

**Supplementary Figure 6. Biomarker analysis from RORγt antagonist treated tumors**. qPCR expression analysis of indicated genes in vehicle, Cmpd 3 or Cmpd 3 plus Gemcitabine treated tumor samples. Replicate values plotted individually with mean ± SD represented.

## Discussion

There is a multitude of pre-clinical data demonstrating that targeting the RORγt/IL-17A/IL-23 pathway ameliorates disease pathology in multiple autoimmune and inflammatory diseases. In addition, recent clinical successes of Otezla (Apremilast) and VTP-43742 (Vitae Pharmaceuticals) in psoriasis, psoriatic arthritis, autoimmune uveitis and/or ankylosing spondylitis through reduction in circulating IL-17A levels [10, 11] support that targeting RORγt is a viable and potentially high value therapeutic strategy for IL-17-driven autoimmune disorders. Herein we report the design, synthesis and pre-clinical characterization of 3 potent, selective allosteric RORγt inhibitors (Cmpds 1, 2 and 3) with structural similarity but also notable differences to those previously reported by Lycera and Merck [18, 19]. Cmpd 3 emerged as a lead candidate based upon its favorable potency and selectivity, as well as its cross-species pharmacokinetic profile, synthetic accessibility and physicochemical properties. With a quality lead molecule in hand, we set out to interrogate the impact of allosteric inhibition of RORγt across models of immunology and oncology *in vitro* and *in vivo* systems.

RORγt is the central transcriptional regulator of γδT cells, group 3 innate lymphoid cell (ILC3), differentiating Th17 cells and memory Th17 cells [22, 23], promoting expression of key subset effectors, including the lineage defining cytokine IL-17A. Previous reports have demonstrated that RORγt-selective small molecule inhibitors can potently block pro-inflammatory IL-17A cytokine production in differentiating Th17 cells as well as in memory Th17 cells, whose expression of RORγt is pre-existing [26-29]. Consistent with these reports, the RORγt inhibitors Cmpd 1, Cmpd 2 and Cmpd 3 ablated Th17 cell differentiation and Th17 maintenance in human primary cells *in vitro*. Blockade of IL-17A secretion was achieved in a dose dependent manner with single digit nM IC_50_ concentrations, with no overt cell cytotoxicity. Interestingly, ∼95% maximal inhibition with respect to Th17 differentiation corresponded to only ∼60% inhibition in a Th17 maintenance assay. These data suggest that, within memory Th17 cells with pre-existing RORγt expression, a substantial fraction of the IL-17A production is independent of RORγt activity. As expected, the RORγt antagonists did not have a significant impact on the frequency of cells producing the Th1 cell hallmark cytokine IFNγ, indicative of T helper lineage specificity.

In addition to potently inhibiting IL-17A responses in Th17 cells *in vitro*, we have demonstrated that allosteric RORγt inhibitors can ameliorate RORγt-dependent inflammation *in vivo*. Systemic administration of Cmpd 3 significantly reduced IMQ-induced ear thickening and Th17-cytokine gene expression. Further, treatment with Cmpd 3 significantly attenuated EAE severity, delayed disease onset and led to significant reductions in body weight loss that were maintained for the duration of study. Moreover, plasma concentrations of Cmpd 3 and *Il17a* or *Il17f* expression levels were inversely correlated with inhibition of Th17-associated gene expression. Importantly, attenuated IMQ-induced skin inflammation & EAE disease pathogenesis was only observed at doses that achieved IC_50_ coverage in excess of 18 hours, suggesting that extended time over IC_50_ is required for a durable response. Small molecule inhibition of RORγt resulted in ∼75% inhibition of RORγt-dependent inflammation could be observed in both acute and chronic inflammatory model systems.

Pancreatic ductal adenocarcinoma accounts for ∼95% of all pancreatic cancer cases for which the 5-year survival rate is only 8%. Surgical resection offers the best chance for increased survival, but only 20% of patients are diagnosed early enough to be candidates. Therefore, discovery of novel drivers and treatments for PDAC remains a high unmet need. A recent finding highlighted a role for RORγt for PDAC tumor growth and CSC maintenance [16]. RORγt antagonists, which are currently under investigation in autoimmune and inflammatory diseases, could therefore represent a novel treatment mechanism for PDAC. Our current study investigated the action of the RORγt antagonist Cmpd 3 in both *in vitro* and *in vivo* PDAC models. As expected, Cmpd 3 demonstrated greater inhibition of organoid formation compared to proliferation consistent with the proposed role for RORγt in CSC growth. Inverse agonist SR2211 demonstrated greater inhibition of organoid growth and formation suggesting that mode of inhibition may effect inhibitory potential. However off-target effects of SR2211 cannot be ruled out as differential activity was observed both *in vitro* and *in vivo*. Additionally, while Cmpd 3 decreased KP/C tumor growth, it was not statistically significant compared to vehicle treatment and did not provide an additive effect in combination with gemcitabine. RORγt antagonists currently under investigation in autoimmune and inflammatory diseases may represent a novel treatment option for PDAC but further work is needed to determine a precise mechanism of action and predict response to various inhibitors.

In this report we detailed the pre-clinical characterization of 3 selective and potent allosteric RORγt inhibitors and demonstrated inhibition of RORγt activity and subsequent RORγt-dependent inflammatory responses in multiple immune cells both *in vitro* and *in vivo*. A maximum of ∼75% inhibition of RORγt-dependent inflammation was achieved in acute and chronic inflammatory settings. Interestingly, in our hands, VTP-43742, required extended IC_90_ coverage to achieve ∼45% inhibition of Th17-dependent responses whereas Cmpd 3 achieved the same level of response with IC_50_ coverage alone. Given the clinical impact of VTP-43742 [11] and the proposed advantage of allosteric inhibition, Cmpd 3 may represent a complimentary treatment option for psoriasis.

There is a breadth of literature supporting a therapeutic benefit for targeting RORγt in Th17-driven autoimmune indications, however, there is also a potential safety liability in that knockout of RORγt in adult mice leads to development of lymphoblastic lymphomas within 6 months, in a manner similar to embryonic RORγt loss [14, 37]. Multiple reports have identified RORγt inhibitors as causative agents in inducing lymphomas in rodents and non-human primates [33, 34]. Further, this effect is thought to be driven by RORγt mediated modulation of *Bclxl* expression in double negative (DN) T cell populations in the thymus, a phenomenon we observed in our studies. Human subjects with RORC knockout have been identified and do not exhibit signs of lymphoma [38], however, given our observations and additional reports of RORγt inhibition-induced thymocyte apoptosis in rodents & non-human primates (unpublished results: Bristol Myers Squibb (Haggerty et al. Society of Toxicology conference, 2020) and Genentech (Zbieg et al. Federation of Clinical Immunology Societies conference, 2018) the question remains whether the susceptibility to thymic lymphomas is a rodent-specific phenomenon and whether this presents a significant safety liability in human.

While the risk of lymphoma represents a significant hurdle for the development of RORγt inhibitors for the treatment of chronic autoimmune diseases, novel allosteric RORγt inhibitors reported herein may serve as additional tools for interrogation of RORγt biology. Decoupling of RORγt pathway inhibition and risk of thymic lymphomas, through suppression of Th17-mediated pathology, would represent a breakthrough with respect to the use of RORγt inhibitors for the treatment of autoimmune disorders, and would have the potential to provide a significant advancement in treatment options for patients worldwide.

## Author Contributions

All authors designed, performed experiments and/or analyzed data. JME and LE led the research. SAS, AL, LE and JME conceptualized and wrote the manuscript.

## Acknowledgements

We would like to thank all current and former employees of Celgene Corp and Bristol Myers Squibb that contributed to this work.

